# Unintended inhibition of protein function using GFP nanobodies in human cells

**DOI:** 10.1101/590984

**Authors:** Cansu Küey, Gabrielle Larocque, Nicholas I. Clarke, Stephen J. Royle

## Abstract

Tagging a protein-of-interest with GFP using genome editing is a popular approach to study protein function in cell and developmental biology. To avoid re-engineering cell lines or organisms in order to introduce additional tags, functionalized nanobodies that bind GFP can be used to extend the functionality of the GFP tag. We developed functionalized nanobodies, which we termed “dongles”, that could add, for example, an FKBP tag to a GFP-tagged protein-of-interest; enabling knocksideways experiments in GFP knock-in cell lines. The power of knocksideways is that it allows investigators to rapidly switch the protein from an active to an inactive state. We show that dongles allow for effective knocksideways of GFP-tagged proteins in genome-edited human cells. However, we discovered that nanobody binding to dynamin-2-GFP caused inhibition of dynamin function prior to knocksideways. While this limitation might be specific to the protein studied, it was significant enough to convince us not to pursue development of dongle technology further.

## Introduction

Fluorescent proteins revolutionized cell biology. The green fluorescent protein (GFP) or its relatives can be attached to virtually any protein-of-interest and allow the direct visualization of that protein by light microscopy (Wang and Hazelrigg, 1994). Whole genome GFP-tagging projects have been completed in yeast (Huh et al., 2003), plants (Tian et al., 2004), bacteria (Kitagawa et al., 2005), and fly (Nagarkar-Jaiswal et al., 2015). The advent of genome engineering, particularly via CRISPR/Cas9, has allowed the creation of GFP knock-in mammalian cell lines in labs around the world (Jinek et al., 2013), with centralized efforts to systematically tag genes in human induced pluripotent stem cells (Roberts et al., 2017). While these resources are incredibly useful, additional tags would further enhance our ability to probe protein function in single cells.

Of particular interest is the ability to rapidly modulate protein function. Inducible methods such as relocation (Haruki et al., 2008; Robinson et al., 2010) and degradation (Nishimura et al., 2009) allow investigators to study the effect of inactivating a protein-of-interest in live cells. For example, we have used the knocksideways method to study protein function at distinct stages of mitosis, without perturbing interphase function (Cheeseman et al., 2013). Here, a protein-of-interest has an FKBP tag that allows inducible binding to a mitochondrially targeted protein containing an FRB tag (Mito-Trap) via the heterodimerization of FKBP and FRB by rapamycin (Robinson et al., 2010). The power of these methods lies in the comparison of the active and inactive states of the protein-of-interest.

The development of camelid nanobodies that bind GFP have been very useful as affinity purification tools (Roth-bauer et al., 2008). Since these nanobodies can be readily expressed in cells, it is possible to use them as “dongles”: to extend the functionality of GFP by attaching a new protein domain to the GFP-tagged protein-of-interest via fusion with the nanobody. This approach has been exploited to degrade proteins-of-interest (Caussinus et al., 2011; Kanner et al., 2017; Daniel et al., 2018; Yamaguchi et al., 2019), to introduce additional tags (Rothbauer et al., 2008; Ariotti et al., 2015; Derivery et al., 2017; Zhao et al., 2018), or to constitutively relocalize GFP-tagged proteins (Schornack et al., 2009; Derivery et al., 2015). Recently a suite of functionalized nanobodies to GFP or RFP were generated enabling recoloring, inactivation, ectopic recruitment, and calcium sensing (Prole and Taylor, 2019). The dongle approach holds much promise because it is flexible and saves investigators from re-engineering knock-in cell lines to introduce additional tags.

Some time ago, we developed dongles to allow knock-sideways experiments in GFP knock-in cell lines. The approach certainly works and we demonstrate this using two different genome-edited human cell lines. However, we discovered during the course of development that nanobody binding to dynamin-2-GFP causes inhibition of dynamin function, prior to any induced inactivation. Since the purpose of knocksideways is to compare active and inactive states, the dongles could not be used in this way. The aim of this paper is to alert other labs to the possibility that nanobodies against GFP can cause inhibition of function of the GFP-tagged target protein. While this limitation might be restricted to dynamin, it was significant enough to convince us that we should not pursue dongles further as a cell biological tool. We discuss what strategies investigators might pursue as alternatives and outline possible applications of dongles despite this limitation.

## Results

### Testing fluorescent protein selectivity of dongles in cells

Most experimental applications of “dongles” would involve two different fluorescent proteins, one as a target for the dongle and a second as an experimental readout. We therefore wanted to assess the fluorescent protein selectivity of the GFP nanobody in cells. To do this, we used a visual screening method in HeLa cells by expressing GFP nanobody (GFP-binding protein-enhancer, GBPen) that was constitutively attached to the mitochondria (DongleTrap, see Methods) along with a suite of twenty-five different fluorescent proteins. Affinity of the fluorescent protein for the DongleTrap resulted in a steady-state relocation to the mitochondria, while lack of interaction meant that the fluorescent protein remained cytoplasmic (Figure 1). We found relocation for mAzurite, EBFP2, sfGFP, mEmerald, EGFP, Clover, EYFP, mVenus, and mCitrine. While the following fluorescent proteins remained cytoplasmic in all cells examined: TagBFP2, ECFP, mCerulean3, mTurquoise2, mAzamiGreen, mNeonGreen, mOrange2, mKO2, DsRed, mRuby2, mScarlet, mRFP, mCherry, mNeptune2, mMaroon, and TagRFP657. All of the fluorescent proteins that DongleTrap binds are derivatives of avGFP (GFP from *Aequorea victoria*), while it did not bind proteins from other lineages, e.g. dsRed, eqFP578, and LanYFP (Lambert, 2018). The GBPen has further specificity besides lineage, since DongleTrap did not bind other avGFP descendants ECFP, mCerulean3 and mTurquoise2 (Kubala et al., 2010). These experiments demonstrated which tags could be manipulated by dongles in cells (e.g. GFP), but also which fluorescent proteins can be used simultaneously with these tools, without interference (e.g. mCherry).

**Figure 1.**
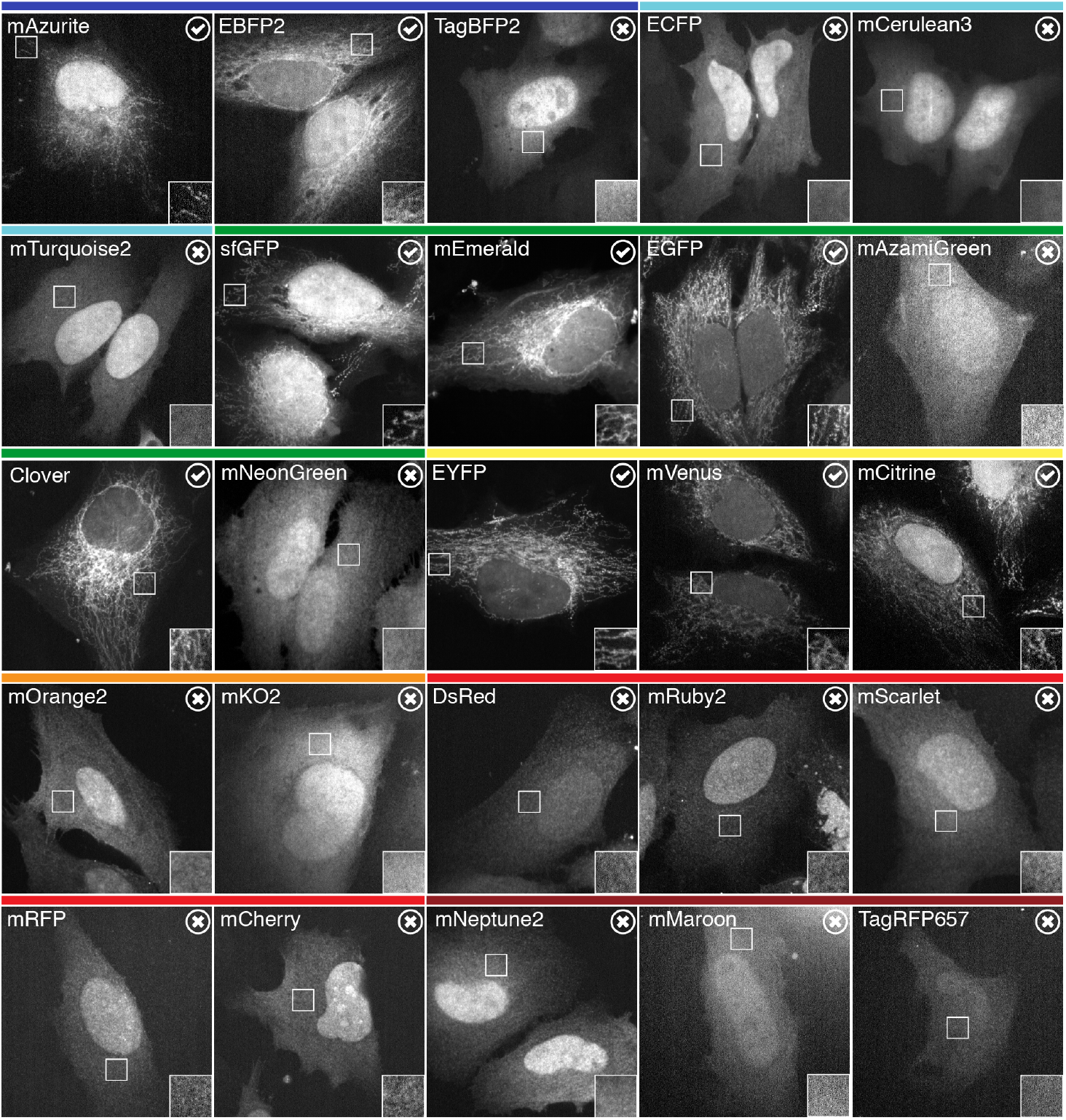
Selectivity of dongles for fluorescent proteins. Representative images of HeLa cells expressing DongleTrap (pMito-GBPen) and the indicated fluorescent protein. Binding to DongleTrap results in mitochondrial localization of the fluorescent protein and is indicated by a tick/check mark. Fluorescent proteins that do not bind to the DongleTrap remain cytoplasmic and are indicated by a cross. Insets show a 2× zoom of the indicated ROI. Colored bars above indicate the approximate emission of the fluorescent proteins tested.

### Dongles can be used to extend the function of GFP

Knocksideways is a useful tool to rapidly inactivate proteins by sequestering them on to mitochondria using heterodimerization of FKBP and FRB domains (Robinson et al., 2010). Typically, the FKBP domain is fused to the protein-of-interest (usually along with GFP for visualization) and the FRB domain is part of MitoTrap (Figure 2). To demonstrate the usual application of this method, we rerouted the membrane trafficking protein Tumor Protein D54 (TPD54/TPD52L2) to mitochondria (for detailed analysis of TPD54 rerouting see Larocque et al., 2018). To do this, we expressed GFP-FKBP-TPD54 in HeLa cells either alone or together with MitoTrap. Application of 200 nM rapamycin caused the relocation of GFP-FKBP-TPD54 to mitochondria in seconds, only when MitoTrap was present (Figure 2A).

**Figure 2.**
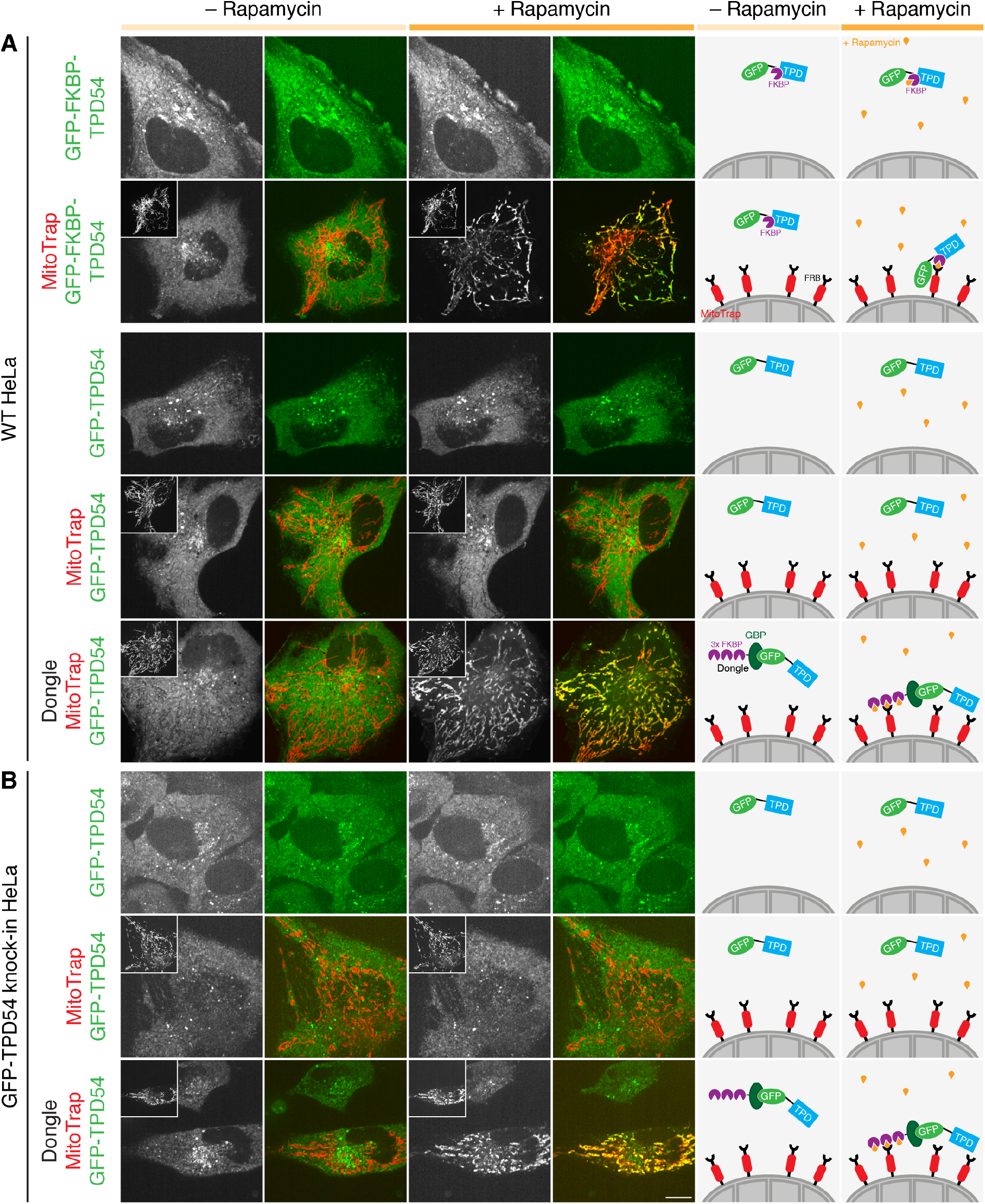
Dongles allow for knocksideways of GFP-tagged proteins that have no FKBP tag. Representative confocal images of live cells taken before (light orange) or after (dark orange) addition of 200 nM rapamycin. **A**, GFP-FKBP-TPD54 or GFP-TPD54 was expressed in WT HeLa cells, along with MitoTrap (Mito-mCherry-FRB) alone or together with the dongle as indicated. If MitoTrap (red in merge) was co-expressed, the red channel is shown inset at half-size. **B**, In GFP-TPD54 knock-in HeLa cells, MitoTrap or MitoTrap + dongle was expressed as indicated. Scale bar, 10 µm. Schematic diagrams to the right illustrate the experimental conditions and the respective result.

We wanted to use knocksideways on proteins that have a GFP-tag, but no FKBP. To enable this we designed a dongle comprising three copies of FKBP fused to the N-terminus of GBPen, which can be co-expressed in cells along with MitoTrap (see Methods). When expressed transiently in cells along with GFP-TPD54, application of rapamycin (200 nM) caused GFP-TPD54 to become rapidly rerouted to mitochondria (Figure 2A). Mitochondrial rerouting was dependent on the presence of the dongle, since no rerouting was seen in rapamycin-treated cells expressing GFP-TPD54 and MitoTrap. The effect was indistinguishable from rerouting of GFP-FKBP-TPD54 to MitoTrap in response to rapamycin (Figure 2A).

Given these encouraging results, we next tested if dongles could be used to reroute endogenous proteins, tagged with GFP, to the mitochondria. To do this, we expressed the dongle and MitoTrap in cells where endogenous TPD54 was tagged with GFP (Larocque et al., 2018). We found that GFP-TPD54 was rerouted when the dongle and MitoTrap were present and rapamycin was added (Figure 2B). Knocksideways was qualitatively similar to GFP-FKBP-TPD54 or GFP-TPD54 and dongle, expressed with MitoTrap in wild-type HeLa cells (Figure 2). These experiments indicate that the dongles can be used to extend the function of GFP and to permit knocksideways experiments in GFP knock-in cell lines without an FKBP tag. We termed this method “dongle-knocksideways”.

### Knocksideways of dynamin-2 in gene-edited human cells

We next wanted to use the dongle-knocksideways method to switch-off endocytosis on-demand. A direct approach would be to inactivate the large GTPase dynamin, which is essential for vesicle scission during endocytosis (Antonny et al., 2016). We therefore tested dongle-knocksideways in SK-MEL-2 hDNM2^EN-all^ cells, where both alleles of dynamin-2 are tagged with GFP (Doyon et al., 2011). Confocal imaging revealed rapid and efficient rerouting of dynamin-2-GFP (DNM2-GFP) to mitochondria using 200 nM rapamycin in cells co-expressing the dongle and MitoTrap (Figure 3A and Supplementary Video SV1).

**Figure 3.**
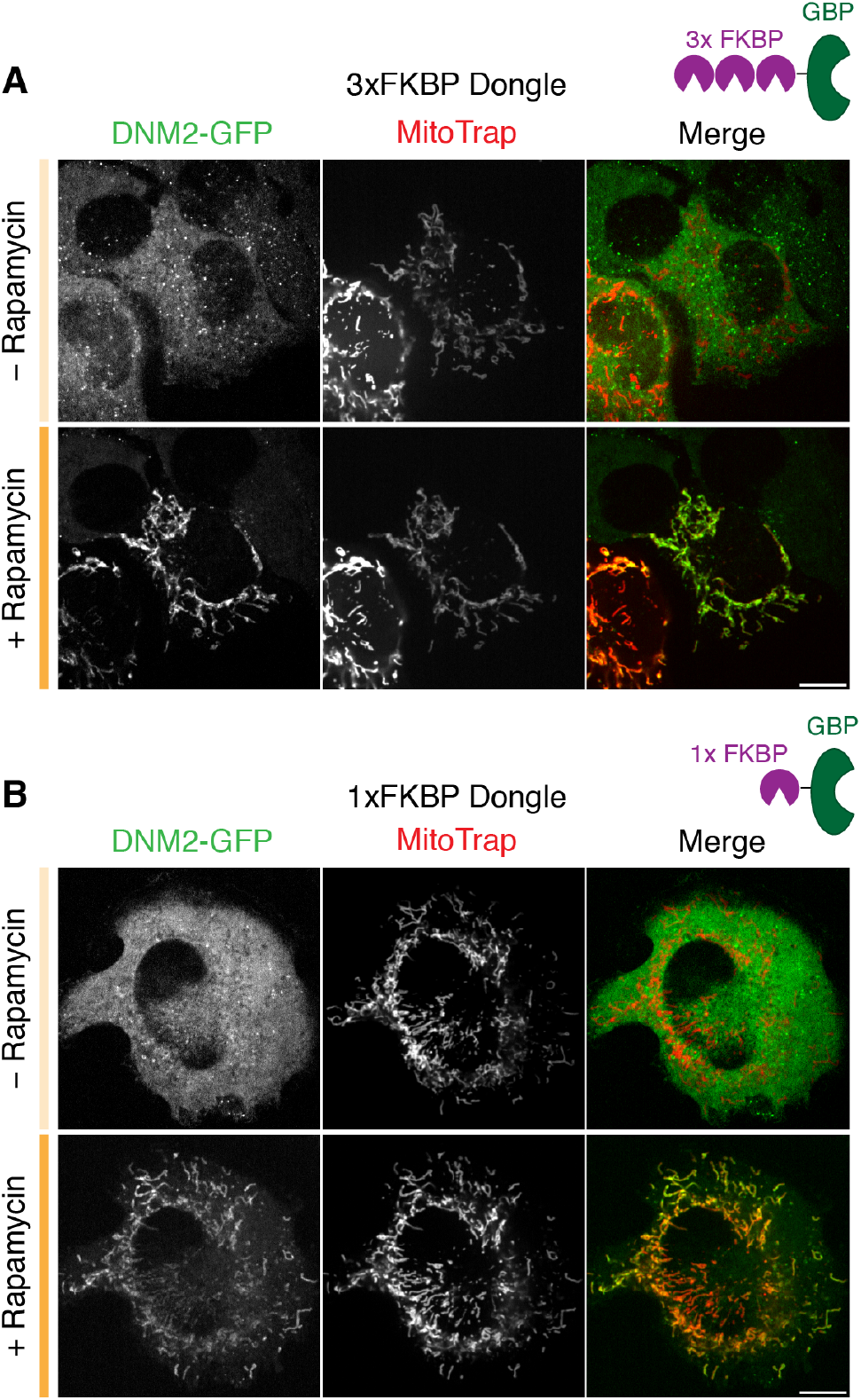
Dongle-knocksideways efficiently reroutes dynamin-2-GFP to mitochondria. Still confocal images from dongle-knocksideways experiments showing a cell before and after application of 200 nM rapamycin. SK-MEL-2 hDNM2^EN-all^ cells expressing MitoTrap and either 3xFKBP Dongle (**A**) or 1xFKBP Dongle (**B**). Scale bars, 10 µm.

The dongle used for knocksideways has three FKBP domains in tandem, attached to GBPen (3xFKBP Dongle). For reasons that will become clear below, we also generated a dongle with a single FKBP domain (1xFKBP Dongle). Using this construct for dongle-knocksideways of DNM2-GFP in SK-MEL-2 hDNM2^EN-all^ cells was similar to experiments that used 3xFKBP Dongle (Figure 3B). Therefore, dynamin-2-GFP can be rerouted efficiently to mitochondria using dongle-knocksideways.

### Inhibition of clathrin-mediated endocytosis using dongles

Does dongle-knocksideways of dynamin-2-GFP cause an inhibition of clathrin-mediated endocytosis (CME)? To answer this question we analyzed the cellular uptake of fluorescently-conjugated transferrin, an established assay for CME. We tested the effect of dongle-knocksideways (rapamycin *versus* vehicle) and compared this to inhibition of endocytosis (sucrose) using hypertonic media (Hansen et al., 1993). To our surprise, we found that expression of the dongle was sufficient to inhibit CME in SK-MEL-2 hDNM2^EN-all^ cells (Figure 4). The amount of transferrin uptake in cells expressing MitoTrap together with 3xFKBP Dongle was significantly reduced compared to untransfected cells or those expressing MitoTrap alone (Figure 4). This unintended inhibition of CME was similar to treatment with hypertonic media, a classical method to inhibit endocytosis. We wondered whether the size of 3xFKBP Dongle caused this inhibition and so we generated a 1xFKBP Dongle, which was approximately half the size (3xFKBP Dongle, 49.8 kDa; 1xFKBP Dongle, 25.9 kDa), and verified that the 1xFKBP Dongle was fully functional for rerouting experiments (Figure 3B). Again, this dongle caused inhibition of CME by expression in SK-MEL-2 hDNM2^EN-all^ cells, similar to that seen for 3xFKBP Dongle (Figure 4). Similar results were seen using SK-MEL-2 hCLTA^EN^/hDNM2^EN^ cells, which indicated that this effect was not specific to the hDNM2^EN-all^ cell line used (Figure S1). Note that, with either dongle and either cell line, no further inhibition of CME was observed by sucrose treatment nor by rapamycin addition causing dongle-knocksideways. These observations mean that the dongle method cannot be used in this way to inhibit endocytosis on-demand, since the active state is inhibited unintentionally.

**Figure 4.**
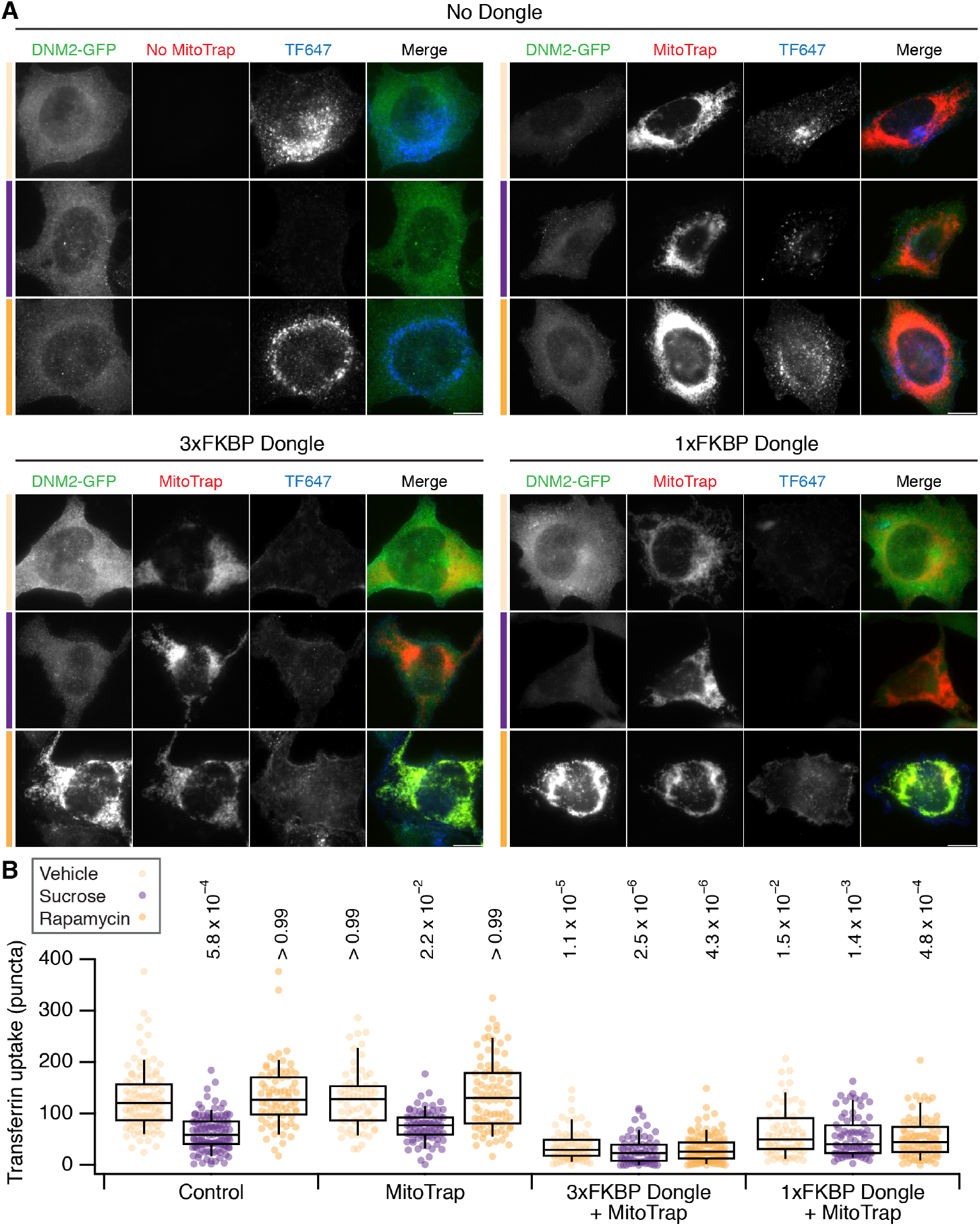
Effect of dongle expression on transferrin uptake in SK-MEL-2 hDNM2^EN-all^ cells. **A**, Micrographs of SK-MEL-2 hDNM2^EN-all^ cells treated with vehicle (light orange), sucrose (purple) or 200 nM rapamycin (orange). Cells were untransfected (No Dongle, No Mitotrap), or expressed MitoTrap alone, or MitoTrap with 3xFKBP Dongle or 1xFKBP Dongle. DNM2-GFP (green), MitoTrap (red) and transferrin-Alexa647 (blue) are displayed using the same minimum and maximum value per channel for all images in the figure. Scale bars, 10 µm. **B**, Box plot to show quantification of transferrin uptake. Expression and treatments are as indicated and colored as in A. Dots represent individual cells from multiple experiments. Box represents the IQR, the line the median and the whiskers the 9^th^ and 91^st^ percentile. *n*_cell_ = 64 - 114, *n*_exp_ = 7. Two-way ANOVA on experimental means, within subject, Factor A = expression, *DF* = 2, *F* = 12.44, *F*_*c*_ = 8.77, *p* < 0.001; Factor B = treatment, *DF* = 3, *F* = 35.02, *F*_*c*_ = 7.05, *p* < 0.001; *A* × *B,DF* = 6, *F* = 3.17, *F*_*c*_ = 5.12, *p* > 0.001. P-values from Dunnett’s *post hoc* test are shown (Control-Vehicle as the control group).

Our results suggested that the unintentional inhibition of CME is caused by dongles binding to dynamin-2-GFP and inhibiting its function. An alternative hypothesis is that the dongles inhibit CME via some unknown mechanism and the effect in cells with dynamin-2-GFP was coincidental. To test if dongles inhibited CME directly, we measured transferrin uptake in HeLa cells with no dynamin-2-GFP, that expressed either the 3xFKBP or 1xFKBP Dongles with MitoTrap (Figure 5). We found that transferrin uptake in these cells was similar to cells expressing MitoTrap alone or to untransfected controls (Figure 5). CME in cells expressing either dongle could be inhibited by sucrose and not by rapamycin treatment which is to be expected if there is no direct inhibition of CME caused by the dongle. These experiments ruled out an inhibitory effect of dongles on CME, and implicate the inhibition seen in cells expressing dynamin-2-GFP as being due to binding dynamin-2-GFP with the nanobody. In conclusion, while the dongles work as a way to reroute proteins to the mitochondria, the inhibition of protein function prior to rapamycin application means that they might not be useful for switching between active and inactive states.

**Figure 5.**
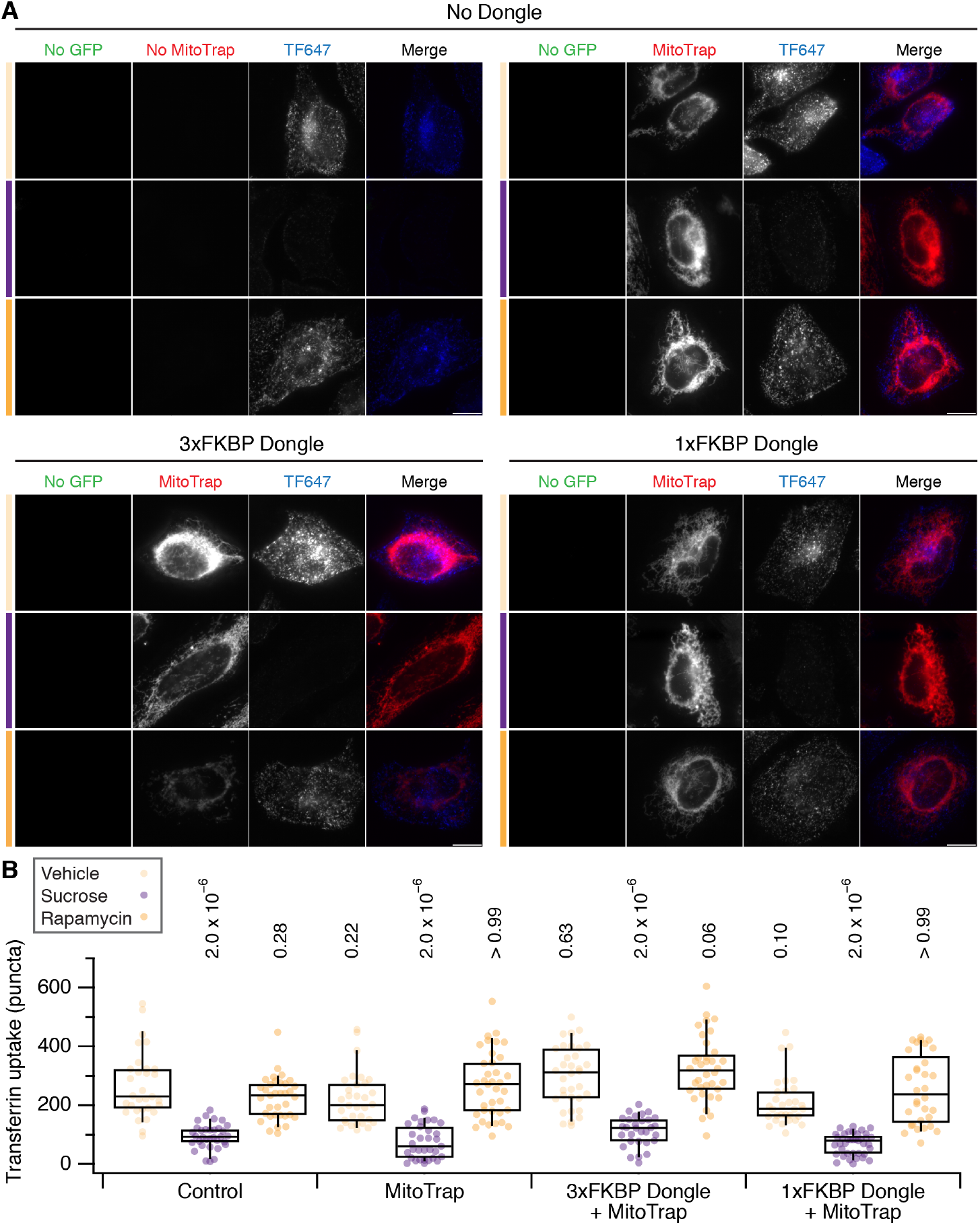
Effect of dongle expression on transferrin uptake in HeLa cells. **A**, Micrographs of HeLa cells treated with vehicle (light orange), sucrose (purple) or 200 nM rapamycin (orange). Cells were untransfected (No Dongle, No Mitotrap), or expressed MitoTrap alone, or MitoTrap with 3xFKBP Dongle or 1xFKBP Dongle. No GFP (green), MitoTrap (red) and transferrin-Alexa647 (blue) are displayed using the same minimum and maximum value per channel for all images in the figure. Scale bars, 10 µm. **B**, Box plot to show quantification of transferrin uptake. Expression and treatments are as indicated and colored as in A. Dots represent individual cells from a single experiment. Box represents the IQR, the line the median and the whiskers the 9^th^ and 91^st^ percentile. P-values from Dunnett’s *post hoc* test are shown (Control-Vehicle as the control group).

## Discussion

In this paper, we described the development of dongles to extend the functionality of GFP for knocksideways experiments. We found that these molecular tools were effective at binding GFP and permitting the rerouting of the target protein to mitochondria. However, we discovered an unintended side-effect: dongles can inhibit the function of the GFP-tagged protein under study.

The use of GFP-binding protein as an intracellular tool to manipulate protein function is becoming widespread (Prole and Taylor, 2019; Daniel et al., 2018; Ariotti et al., 2015). The appeal of the method is that proteins tagged with GFP at their endogenous loci can be adapted using dongles to enable inactivation, relocalization, recoloring or other function. This means that existing cell lines or organisms can be “retrofitted” for additional functionality using such tools. Indeed our initial experiments using dongles were very encouraging: the dongles expressed well, we detected no obvious perturbation of subcellular distribution of the target proteins we examined, and they permitted knocksideways which, in the case of TPD54, was very similar to rerouting the same protein with a fused GFP-FKBP tag. However, we found that when the dongles were expressed in cells with both copies of dynamin-2 tagged with GFP, endocytosis was inhibited. This unintended effect seemed to be due to direct inhibition of dynamin function following binding of dynamin-2-GFP by the nanobody (GBPen) portion of the dongle. It is possible that the inhibitory effect we have observed is specific to dynamin-2-GFP. Dynamins may be uniquely sensitive because they self-oligomerize, and we know that their action can be readily inhibited by simple expression of a GTPase deficient isoform (Damke et al., 1994). However, it is likely that dongles inhibit other GFP-tagged proteins and this has deterred us from pursuing this method further.

We were fortunate to test this method on dynamin-2 which has a clear functional readout because it controls the terminal step in clathrin-mediated endocytosis (Antonny et al., 2016). Other proteins do not have such unambiguous readouts, or their function can only be measured indirectly, if at all. This would mean that if dongles were used with these proteins, inhibition would not be revealed and potentially misleading conclusions drawn. This problem is compounded because dongles would be most useful when applied to proteins whose function is uncertain; so the inhibition of protein function caused by these tools may be hidden from the investigator. Our advice is that the same caution and functional tests should be applied when using GFP nanobodies in cells as when generating GFP-tagged proteins themselves (Snapp, 2005). Even so, inhibition may not be revealed in pilot experiments where the GFP-tagged target protein is overexpressed together with the dongle; since endogenous untagged protein or excess GFP-tagged protein which is not bound to the dongle, could substitute functionally for the inhibited protein.

### Mechanism of unintended inhibition

The mechanism of inhibition of dynamin-2-GFP function by dongles is unclear. The simplest explanation is that extending the GFP tag using a GFP nanobody plus additional domains results in a modification that is simply too large for dynamin to function normally; whether this is because of reduced dynamics, blocked interactions or some other mechanism. We saw similar unintended inhibition when the size of the dongle was reduced by half (from three FKBP domains to one) suggesting that the binding of the nanobody itself is inhibitory rather than there being a size limit to the tag that dynamin-2 can tolerate. It is considered that most proteins can tolerate the addition of GFP or GFP-FKBP tag, but it is perhaps underappreciated that at some level, tags will always interfere with protein function. The fact that we see inhibition using dongles is perhaps not surprising.

### Future usage of nanobody-based methods in cells

Unintended inhibition affects experiments where the protein-of-interest needs to be functional (active) prior to inactivation. For other applications of dongles, inhibition may not be such a concern. First, in constitutive mislocalization experiments, where the goal is to chronically inactivate protein function by changing its cellular localization, dongles remain an important tool. Second, it is unclear if labelling strategies based on dongles are compromised by inhibition (Ariotti et al., 2015). We saw no evidence of gross changes in subcellular localization of the two proteins we tested, however, it remains questionable whether imaging a protein-of-interest in its inhibited state is representative of its normal distribution. Third, in cases where investigators simply want to put a functional domain to a new location using a GFP-tagged anchor protein, such as calcium sensors at the endoplasmic reticulum (Prole and Taylor, 2019), inhibition of the anchor may not be a concern. Fourth, our finding of nanobody-mediated inactivation of protein function may even be useful as a general purpose method for inhibiting protein function in gene-edited cell lines.

### Possible approaches to retrofit knock-in cell lines to confer new functions and avoid unintended inhibition

What is the best strategy to extend the functionality of tags introduced with knock-in technology? First, it may be possible to reduce the inhibitory effect of dongles by mutating the GBP moiety or using different domain configurations and/or by changing the linker regions. Second, alternative GFP-binding proteins such as those based on a designed ankyrin repeat protein (DARPin) scaffold may be functionalized and used as dongles (Brauchle et al., 2014). It is possible that these reagents do not have the same inhibitory effects. Third, using split-GFP technology, proteins-of-interest could be tagged with GFP11 and then the fluorescence complemented with a GFP1-10 protein (Kamiyama et al., 2016), where GFP1-10 is fused to other domains to extend the functionality. A further advantage of this third method is that the fluorescence of the tagged protein can also be altered during the complementation (Kamiyama et al., 2016). However, a weakness is that this method would not take advantage of existing GFP-tagged collections, and would require new knock-ins to be generated in most cases.

## Conclusion

We embarked on the dongle project in order to avoid making several knock-in cell lines per protein-of-interest. Knock-in of a GFP tag in a single cell line could be extended via dongles in order to fulfil many different functions, rather than separately knocking in GFP, GFP-FKBP, GFP-AID and so on. We have reluctantly concluded that dongles are too problematic for use and we now generate the specific cell lines we need in order to do the experiments we want to do. Our goal with this paper is to make available the information that we have obtained, so that other groups can decide if dongles are worth pursuing as a cell biological tool.

## Methods

### Molecular biology

Construction of plasmids to express GFP-TPD54 and GFP-FKBP-TPD54 and mCherry-MitoTrap (pMito-mCherry-FRB) was described previously (Cheeseman et al., 2013; Larocque et al., 2018). The nanobody cDNA used in this paper, described as GBPen (GFP-binding protein enhancer), was synthesized from published sequences (Kubala et al., 2010; Kirchhofer et al., 2010). To make pMito-mCherry-FRB-IRES-FKBP(III)-GBPen, a bicistronic vector to co-express mCherry-MitoTrap and 3xFKBP-GBPen via an internal ribosome entry site (IRES), a custom insert was made by gene synthesis (GenScript) and inserted into pEGFP-C1 in place of GFP at AgeI and EcoRI. To express DongleTrap, pMito-GBPen was made by amplifying GBPen from a plasmid containing FKBP(III)-GBPen (Forward: cttaggatccggcaCAGGTGCAGCTG, Reverse: ggcctctagaTCAATGGTGATGGTG) cloning into demethylated pMito-mCherry-FRB using BamHI and XbaI. To make pMito-mCherry-FRB-IRES-FKBP(I)-GBPen, the bit of IRES including the HindIII cut site and 1xFKBP was amplified by PCR from pMito-mCherry-FRB-IRES-FKBP(III)-GBPen with addition of a BglII site at the end of the amplified fragment (Forward: GTTCCTCTGGAAGCTTCTTGAAG, Reverse: gcga-gatctTTCCAGTTTTAGAAGCTCCACATC). The product was cut with HindIII and BglII. Same vector was cut with HindIII and BglII resulting in a vector lacking all three FKBPs. The cut PCR product was ligated back into the cut vector.

Plasmids to express fluorescent proteins were either available from previous work: pDsRed-N1, pEGFP-N1, pECFP-N1, pEYFP-N1, pmScarlet-C1, pmRFP-N1, pmCherry-N1, pTagRFP657-N1, psfGFP-N1, pTagBFP2; from Addgene: pEBFP2-N1 (54595), pmAzurite-N1 (54617), pmCerulean3-N1 (54730), pmTurquoise2-N1 (60561), pmVenus-N1 (27793), pmRuby2-N1 (54614), pmNeptune2-N1 (54837), pmOrange2-N1 (54499), pmCitrine2-N1 (54594), mEmerald-N1 (53976), pcDNA3-Clover (40259), pmAzamiGreen-N1 (54798), pmMaroon-N1 (54554), pmKO2-N1 (54625); or from Allele Biotech: pmNeonGreen-N1.

### Cell biology

HeLa cells (HPA/ECACC #93021013) or GFP-TPD54 knock-in HeLa cells (Larocque et al., 2018) were cultured in DMEM + GlutaMAX (Thermo Fisher) supplemented with 10 % fetal bovine serum, and 100 Uml^-1^ penicillin/streptomycin. SK-MEL-2 hDNM2^EN-all^or hDNM2^EN-all^/CLTA^EN^ cells were cultured in Dulbecco’s Modified Eagle’s Medium/Nutrient Mixture F-12 Ham (Sigma-Aldrich) supplemented with 10 % fetal bovine serum, 1 % L-glutamine, 3.5 % sodium bicarbonate and 100 Uml^-1^ penicillin/streptomycin. All cells were kept at 37 °C and 5 % CO_2_. HeLa cells were transfected with 1.2 µg DNA (total) per 3 µL GeneJuice (Merck Millipore) according to the manufacturer’s instructions. SK-MEL-2 cells were transfected with 4.8 µg DNA (total) per 850,000 cells using Neon Transfection System (Thermo Fisher) with 3 pulses of 1500 V, 10 ms. Cells were analyzed 2 d post-transfection.

Transferrin uptake experiments were as described previously (Clarke and Royle, 2018). Briefly, cells were serum-starved for 30 min. They were exposed to 200 nM rapamycin (Alfa Aesar) or 0.1 % ethanol (vehicle) for 10 min, and then incubated with 100 µg/mL Alexa 647conjugated transferrin (Invitrogen) for 10 min. Hypertonic sucrose media (0.45 M) was used to inhibit transferrin uptake. All incubations were in serum-free media at 37 °C with 5 % CO_2_ in a humidified incubator. Cells were then fixed in 3 % PFA/4 % sucrose in PBS and mounted on slides using Mowiol.

### Microscopy

For live-cell imaging of rerouting experiments, cells were grown in 4-well glass-bottom 3.5 cm dishes (Greiner Bio-One) and media exchanged for Leibovitz L-15 CO_2_-independent medium. Rerouting was triggered by addition of 200 nM rapamycin in L-15 media. All cells were imaged at 37 °C on a spinning disc confocal system (Ultraview Vox; PerkinElmer) with a 100 × 1.4 NA oil-immersion objective. Images were captured using an ORCA-R2 digital CCD camera (Hamamatsu) following excitation with 488 nm and 561 nm lasers. Imaging of fixed cells was done on a Nikon Ti-U epiflorescence microscope with 100x oil-immersion objective, CoolSnap MYO camera (Photometrics) using NIS elements software.

### Data analysis

Analysis of transferrin uptake was done as described previously (Wood et al., 2017). Briefly, single cells were outlined manually in Fiji. Vesicular structures were isolated by applying a manual threshold to images in the transferrin channel. Positive structures were counted using “Analyze particles”, with limits of 0.03 − 0.8 µm and circularity of 0.3 − 1.0 All analysis was done with the experimenter blind to the conditions of the experiment.

Figures were made with FIJI or Igor Pro 8 (WaveMetrics), and assembled using Adobe Illustrator. Null hypothesis statistical tests were done as described in the figure legends.

## ACKNOWLEDGEMENTS

We thank David Drubin for the kind gift of genome-edited SK-MEL-2 cell lines. Members of the lab gave constructive criticism, Penny La-Borde made the GFP-TPD54 knock-in HeLa cells and helped with early stages of the dongle project, and Miguel Hernández González critically read the manuscript. We gratefully acknowledge CAMDU (Computing and Advanced Microscopy Unit) for their support and assistance in this work. CK acknowledges funding from the University of Warwick, the EPSRC and BBSRC Centre for Doctoral Training in Synthetic Biology (grant EP/L016494/1).

## AUTHOR CONTRIBUTIONS

CK: Investigation, Methodology, Formal analysis, Writing—review and editing. GL: Investigation, Methodology, Supervision, Writing—review and editing. NIC: Conceptualization, Investigation, Methodology. SJR: Conceptualization, Resources, Supervision, Investigation, Methodology, Writing—original draft, Project administration, Writing—review and editing.

## COMPETING FINANCIAL INTERESTS

The authors declare no conflict of interest.

## Supplementary Information

**Figure S1.**
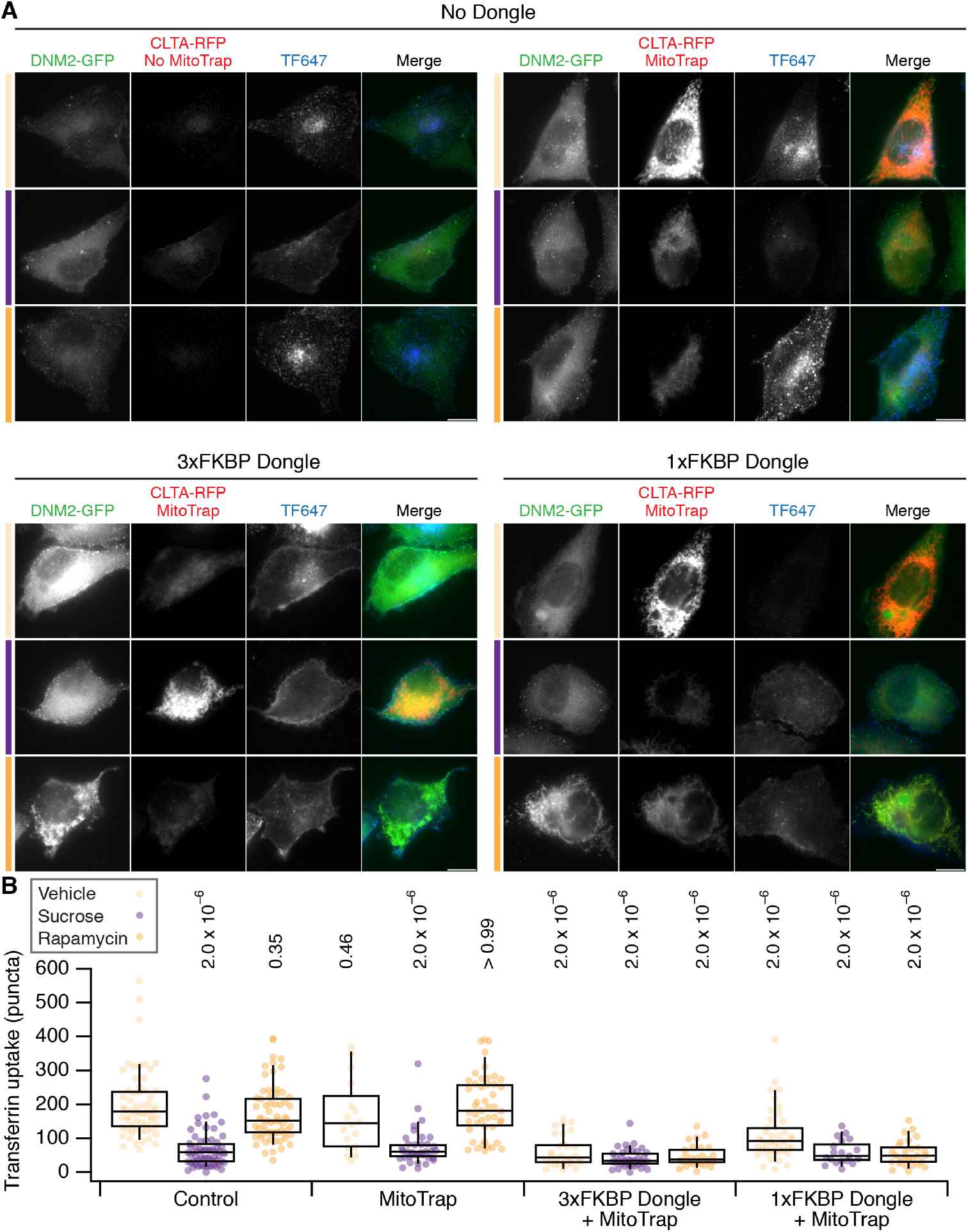
Effect of dongle expression on transferrin uptake in SK-MEL-2 hCLTA^EN^/hDNM2^EN^cells. **A**, Micrographs of SK-MEL-2 hCLTA^EN^/hDNM2^EN^ cells treated with vehicle (light orange), sucrose (purple) or 200 nM rapamycin (orange). Cells were untransfected (No Dongle, No Mitotrap), or expressed MitoTrap alone, or MitoTrap with 3xFKBP Dongle or 1xFKBP Dongle. DNM2-GFP (green), CLTA-RFP + MitoTrap (red) and transferrin-Alexa647 (blue) are shown. Note that MitoTrap and CLTA-RFP appear in the same channel. Scale bars, 10 µm. **B**, Box plot to show quantification of transferrin uptake. Expression and treatments are as indicated and colored as in A. Dots represent individual cells from multiple experiments. Box represents the IQR, the line the median and the whiskers the 9^th^ and 91^st^ percentile. P-values from Dunnett’s *post hoc* test are shown (Control-Vehicle as the control group). *n*_cell_ = 19 - 64, *n*_exp_ = 3.

## Supplementary Videos

**Figure SV1.**
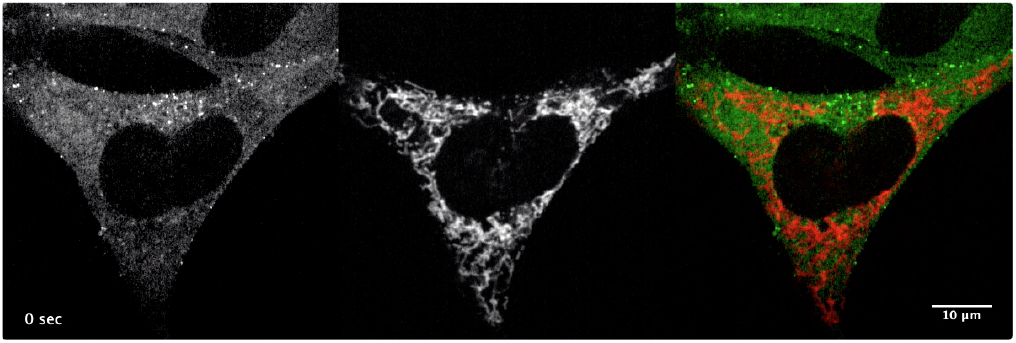
Knocksideways of DNM2-GFP using 3xFKBP Dongle. Live cell confocal microscopy of dongle-knocksideways, 200 nM rapamycin is added at 10 s. Dynamin-2-GFP (left, green) and MitoTrap (middle, red) are shown together with a merge (right). Time, seconds. Scale bar, 10 µm.

## Notes

#### Summary of Updates

- Images in figures where transferrin uptake was quantified are now shown using the same minimum and maximum values per channel, for all images in the figure. - A missing box was added to Figure 1

